# Optimizing Cell Therapy by Sorting Cells with High Extracellular Vesicle Secretion

**DOI:** 10.1101/2023.05.29.542772

**Authors:** Doyeon Koo, Xiao Cheng, Shreya Udani, Dashuai Zhu, Junlang Li, Brian Hall, Natalie Tsubamoto, Shiqi Hu, Jina Ko, Ke Cheng, Dino Di Carlo

## Abstract

Critical challenges remain in clinical translation of extracellular vesicle (EV)-based therapeutics due to the absence of methods to enrich cells with high EV secretion. Current cell sorting methods are limited to surface markers that are uncorrelated to EV secretion or therapeutic potential. We developed a nanovial technology for enrichment of millions of single cells based on EV secretion. This approach was applied to select mesenchymal stem cells (MSCs) with high EV secretion as therapeutic cells for improving treatment. The selected MSCs exhibited distinct transcriptional profiles associated with EV biogenesis and vascular regeneration and maintained high levels of EV secretion after sorting and regrowth. In a mouse model of myocardial infarction, treatment with high-secreting MSCs improved heart functions compared to treatment with low-secreting MSCs. These findings highlight the therapeutic importance of EV secretion in regenerative cell therapies and suggest that selecting cells based on EV secretion could enhance therapeutic efficacy.

## Introduction

Cells produce and secrete numerous bioactive small molecules, proteins, and even larger scale self-assembled structures comprising multiple biomolecules, such as extracellular vesicles (EVs). EVs play critical roles in intercellular communication and have been increasingly recognized, and are being applied, as therapeutic mediators to drive intracellular signaling in target cells to elicit regenerative and anti-inflammatory properties to treat diseases such as osteoarthritis, pulmonary fibrosis, or myocardial damage^1–4^. In fact, EVs have been hypothesized to be key factors driving powerful therapeutic benefits of mesenchymal stem cells (MSCs) when directly introduced *in vivo*^5^. However, there are no methods to select cells based on their production and secretion of EVs, and heterogeneity in the propensity cell therapy products to secrete bioactive compounds like EVs have been highlighted as a potential source for inconsistent therapeutic outcomes^6–8^. Current characterization procedures for EVs and their producing cells use bulk enrichment platforms to isolate secreted EVs from large populations of cells, masking the potential heterogeneity in the secretion phenotype of single cells. No technology has been developed yet to analyze and sort single-cells based on EV secretion levels. We hypothesized that technology that can isolate single cells based on high EV secretion could be used to select base cell populations that would be therapeutically superior. Such an approach could also be used uncover key mediators of biogenesis and secretion pathways for EVs, by combining with transcriptomic assays.

Here, we first developed a nanovial technology to characterize and sort millions of cells based on their secretion of EVs with specific tetraspanin molecular markers (Figure 1) and then applied this approach to select therapeutic cells for treatment. We found that MSCs selected by this phenotype were transcriptionally distinct, expressing markers of EV biogenesis and vascular regeneration, and maintained high levels of EV secretion following sorting and regrowth. We applied the technique to select MSCs based on EV secretion and implanted these cells to treat cardiac injury in a mouse model of myocardial infarction. Animals treated with cells selected based on high EV secretion had improved functional and tissue remodeling outcomes compared to those with low EV secretion. These results support the hypothesis that EV-secretion is a therapeutically important aspect for regenerative cell therapies, and can be selected for and/or engineered in the future to improve therapeutic efficacy and reduce batch-to-batch variability.

**Figure 1.**
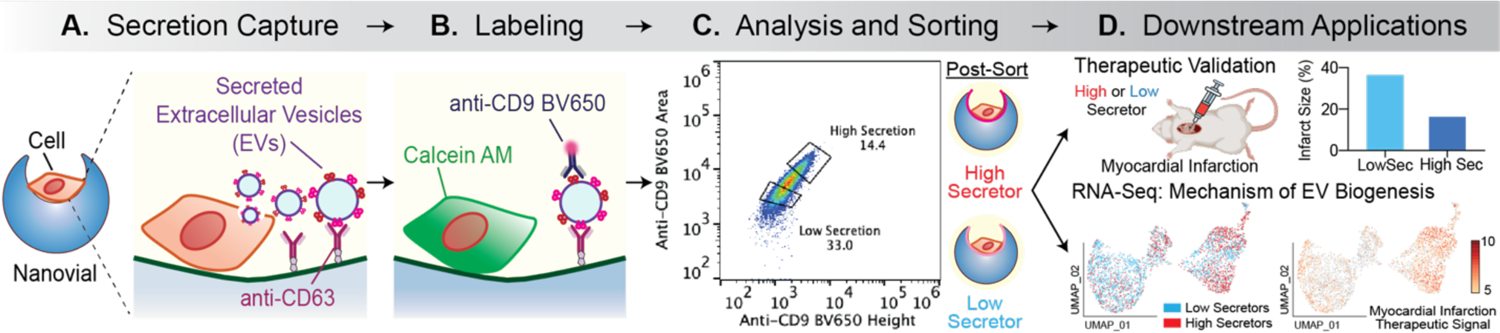
Overview of isolation and downstream analysis of cells based on amount of EV secretion. A) Cells are loaded in nanovials (blue) that are conjugated with anti-CD63 capture antibodies and allowed to secrete EVs. B) Secreted EVs from each cell are captured by the surrounding nanovial and labeled with a fluorescent antibody against CD9 on the surface of EVs. C) Nanovials and corresponding single cells are sorted based on the level of fluorescence signal associated with the captured EVs (High vs. Low Secretors) using a standard cell sorter. D) Downstream transcriptomic analysis is performed on the High or Low Secretor cells to characterize gene expression differences. *In vivo* studies are performed with cells expanded from High Secretor or Low Secretor populations to characterize the effect on therapeutic outcome.

## Results

Since there are no currently validated surface markers that label high-EV secreting cells, we first had to develop a new method to select cells based on their secretion of EVs. Our general strategy made use of cavity-containing hydrogel particles, called nanovials, into which single cells can be seeded. Nanovials were fabricated using a microfluidic device that generates uniform water-in-oil emulsions to create millions of monodisperse polyethylene glycol (PEG)-based nanovials with an inner cavity selectively coated with biotinylated gelatin (Supplementary Figure 1A)^9^. Secreted EVs are captured on the nanovial surface with conjugated antibodies based on expression of unique markers on the secreted EVs (Figure 1A). The captured EVs can then be labeled with fluorescent antibodies against other surface markers (Figure 1B). Finally, the live secreting cells can be sorted using standard fluorescence activated cell sorters based on the level of secretion of EVs for downstream applications (Figure 1C-D).

### MSC binding and capture of CD63+CD9+ EVs on nanovials

We first characterized the marker distribution of the EVs secreted by an immortalized line of human adipose-derived MSCs. EVs are loaded with a diverse range of proteins but EVs released from most cell types express membrane-bound tetraspanins such as CD63, CD9 and CD81 (Figure 2A). To determine the best combination of EV-targeting capture and detection antibodies for our secretion assay and cells, we characterized the bulk EV distributions produced by a culture of immortalized MSCs (iMSCs) using ExoView (Figure 2B). Ultracentrifuged EV populations from iMSC conditioned media contained 30% CD63+CD9+CD81+, 37% CD63+, 12% CD9+ and 4% CD81+ EVs. To avoid capture and detection antibodies competing for the same binding sites on EVs and have higher confidence of EV-specific staining, we targeted CD63+CD9+ EVs using anti-CD63 antibodies to capture EVs and anti-CD9 antibodies for fluorescent labeling of these captured EVs. Strong CD9 signal was observed when EVs isolated from iMSC conditioned medium were incubated with anti-CD63 labeled nanovials, indicating successful capture and detection of CD63+CD9+ EVs (Figure 2C). There was a two orders of magnitude dynamic range in fluorescence signal measured by flow cytometry for detection of CD63+CD9+ EVs as the concentration of EVs was increased (Supplementary Figure 1B).

**Figure 2.**
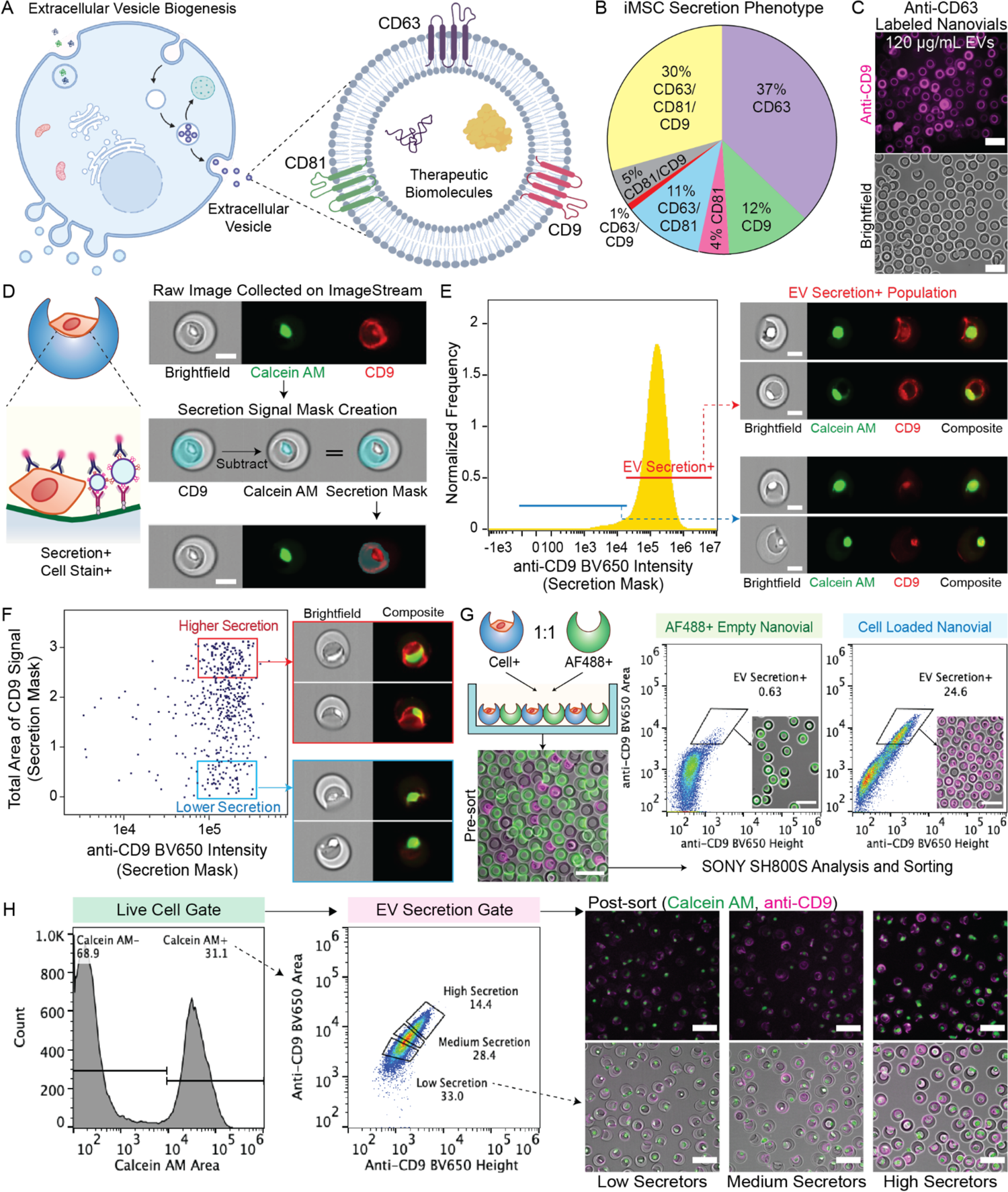
Analysis and sorting of single cells based on EV secretion using nanovials and flow cytometry. A) Schematic of EV biogenesis and expression of tetraspanins (CD63, CD9, CD81) on the surface of EVs. B) Tetraspanin abundance for EVs secreted by immortalized mesenchymal stem cells (iMSC) analyzed by ExoView. C) Validation of sandwich immunoassay for detecting secreted EVs on nanovials. EVs were captured on anti-CD63 labeled nanovials and labeled with fluorescent anti-CD9 antibodies as shown in fluorescence and brightfield microscopic images. Scale bars represent 100 μm. D) Analysis of EV secretion signal on nanovials loaded with cells using ImageStream imaging flow cytometry. A secretion signal mask was created by subtracting the calcein AM signal mask from the CD9 signal mask to exclude non-specific CD9 signal from cell surface staining. Scale bars represent 20 μm. E) EV secretion+ population represents nanovials loaded with cells and containing high anti-CD9 signal spatially located across the cavity from captured EVs. F) Differences in the ratio between the area of secretion signal and overall fluorescence intensity between high and low EV secretors results from spatially distributed secretion signals around the entire surface of the cavity. G) Schematic of an experiment to evaluate cross-talk between cell-loaded and empty nanovials. Less than 1% of empty nanovials were observed to have EV secretion signal when cell-loaded and empty nanovials were maintained in co-culture for 24 hours. H) Single cells on nanovials were sorted based on staining for calcein AM and CD9+ EV secretion signal into three categories of high, medium and low secretors. Fluorescence microscopy images showing sorted populations with corresponding secretion quantity gates. Scale bars represent 100 μm.

### Analysis and sorting based on single-cell EV secretion using nanovials

Our analysis and sorting is focused on single loaded cells and their secreted EVs, so we developed cell loading and FACS gating approaches to enrich this population. Optimum single-cell recovery was possible when a ratio of 1 cell per nanovial was used during seeding in a well plate and nanovials with single cells were further enriched based on fluorescence area vs. width signal of a cytoplasmic dye, Cell Tracker Green (Supplementary Figure 1C). Nanovials with single cells had lower fluorescence width signals as compared to those with two cells (doublets) or aggregates of two nanovials with one or more cells (aggregates), showing successful isolation of these populations (Supplementary Figure 1C).

To characterize the secretion of EVs from cells loaded on nanovials and determine if EV-specific labels could be distinguished from staining of associated cells, we first used imaging flow cytometry (Amnis ImageStream), where staining of both the nanovial surface and cell surface could be independently ascertained. After loading iMSCs onto anti-CD63 labeled nanovials, secreted EVs were accumulated for 24 hours and labeled with fluorescent anti-CD9 antibodies. Due to the presence of some staining with anti-CD9 antibodies on the surface of cells, we created a secretion mask on images to isolate the signal coming from captured CD63+CD9+ EVs on the nanovials rather than signal from cells. The secretion mask comprises a binary mask encompassing the image of the nanovial minus the image of the cell (Figure 2D). Nanovials gated based on higher overall anti-CD9 fluorescence signal in the secretion mask were observed to have anti-CD9 signal associated with the inner surface of the nanovial cavity, while events with low overall intensity reflected signal associated solely with the cell in the nanovial (Figure 2E). Notably, the population of nanovials with high CD9-specific staining across the cavity possessed a greater ratio between the area of staining and overall fluorescence intensity than the population with more localized cell staining, mainly due to the spatially distributed secretion signal that spanned the entire surface of the cavity (Figure 2F). We expected this difference in distribution of fluorescence intensity across the nanovials could also be detected in non-imaging based flow cytometers and FACS instruments by using fluorescence peak shape information.

Considering the higher fluorescence area vs. overall fluorescence intensity we observed for high EV secretors by imaging flow cytometry, we sorted nanovials based on the fluorescence peak area and height values using a standard cell sorter, the SONY SH800S. When setting a secretion signal threshold from negative control nanovials which were not functionalized with any anti-CD63 capture antibody, we were able to identify a population enriched with CD63+CD9+ EV secretion signal. When sorting events in this gate we found it yielded a mixed population of nanovials containing cells with distributed staining across the nanovial surface, reflecting high and low EV secretors (Supplementary Figure 2A). We also observed the highest EV secretion signal when secretion was accumulated over 24 hours compared to 6 or 12 hours of incubation (Supplementary Figure 2B), providing further evidence that the nanovial signal was associated with EV secretion amount.

Given the long incubation time, we were concerned there could be crosstalk of secreted EVs from cells which could be captured by neighboring nanovials. We evaluated crosstalk between nanovials by co-culturing cell-loaded nanovials and fluorescently labeled test nanovials without cells and found that ∼0.6% of test nanovials without cells appeared in the positive secretion gate during the 24-hr incubation period, which compared to ∼25% of cell-loaded nanovials (Figure 2G). Presumably, secreted EVs accumulate at higher concentrations locally, and are localized due to their increased size and reduced diffusivity compared to proteins and the shear-protective cavity of the nanovials.

### Isolation and expansion of high and low EV secretors

Having developed the single-cell EV secretion analysis and sorting workflow, our results suggested that there was significant heterogeneity in the secretion of EVs at the single-cell level. From imaging flow cytometry data, we observed a large distribution of secretion signal area, indicating heterogeneity in the number of EVs each cell secretes (Figure 2F). To better quantify EV secretion at the single-cell level, we simultaneously measured secretion and staining of individual cells with a cell viability dye (calcein AM) (Supplementary Figure 2C). Across the wide distribution of secreting cells, the majority of cells stained positively with calcein AM indicating viability. >90% of calcein AM positive cells on nanovials secreted EV quantities above the detection threshold for negative control nanovials without capture antibodies. However, a high coefficient of variance (CV = 0.49) in the mean fluorescence intensity (6272) indicated a significant dispersion in secretion among EV secretors.

We next wanted to understand whether the propensity for EV secretion of cells sorted using the assay persisted over multiple generations. To investigate the persistence of selected EV secretion phenotypes, we gated and sorted both single cells and groups of 3000 cells based on EV signal (low, medium, high secretors) thresholding above a negative control sample without capture antibody (Figure 2H). Recovery of viable cells with different levels of EV secretion was reflected in the fluorescence microscopy images (Figure 2H). Sorted cells on nanovials expanded in culture actively as cells spread out from the nanovial cavity, adhered to the plate and divided (Supplementary Figure 3A-B). Both clonal populations derived from single MSCs and bulk sorted populations could be expanded over 2 weeks. High secretors from both single-cell and bulk-sorted populations exhibited significantly higher proliferation rates than low secretors (Supplementary Figure 3C). To account for the higher proliferation rate of the expanded populations, an equal number of cells were seeded after 22 and 13 days of expansion for single-cell colony and bulk-sorted populations respectively, and secreted EVs were quantified from conditioned medium. High secretors were observed to maintain a higher EV secretion rate than low secretors from both single-cell and bulk-sorted colonies, indicating that the EV secretion phenotype was maintained even after multiple cell division cycles (Supplementary Figure 3D). The EV production rate from single-cell derived colonies exhibited a larger standard deviation in the number of secreted EVs compared to bulk sorted populations, suggesting the potential for more measurement error in sorting individual cells or the presence of additional heterogeneity at the single-cell level.

### Transcriptomic analysis of low and high EV-secreting cells

We hypothesized that high EV secretors would have differentially expressed genes compared to low EV secretors and specific molecular mechanisms might drive robust EV shedding. We sorted iMSCs loaded on nanovials as high and low EV secretors based on the CD63+CD9+ EV secretion level and profiled single-cell gene expression for these separate populations using the 10X Genomics Chromium system. We visualized the gene expression of both populations using Uniform Manifold Approximation and Projection (UMAP) and overlaid information on whether each cell belonged to the high and low secretor populations. Populations representing each secretion phenotype clustered in different regions of the UMAP. High secretors dominated clusters 1, 2, and 3, while the majority of clusters 4, 5, 6, and 7 contained low secretors (Figure 3A). iMSC stem cell marker expression was consistent across each cluster, indicating that different sub-populations of iMSCs, not differentiation, may be responsible for the observed heterogeneity (Supplementary Figure 4A).

**Figure 3.**
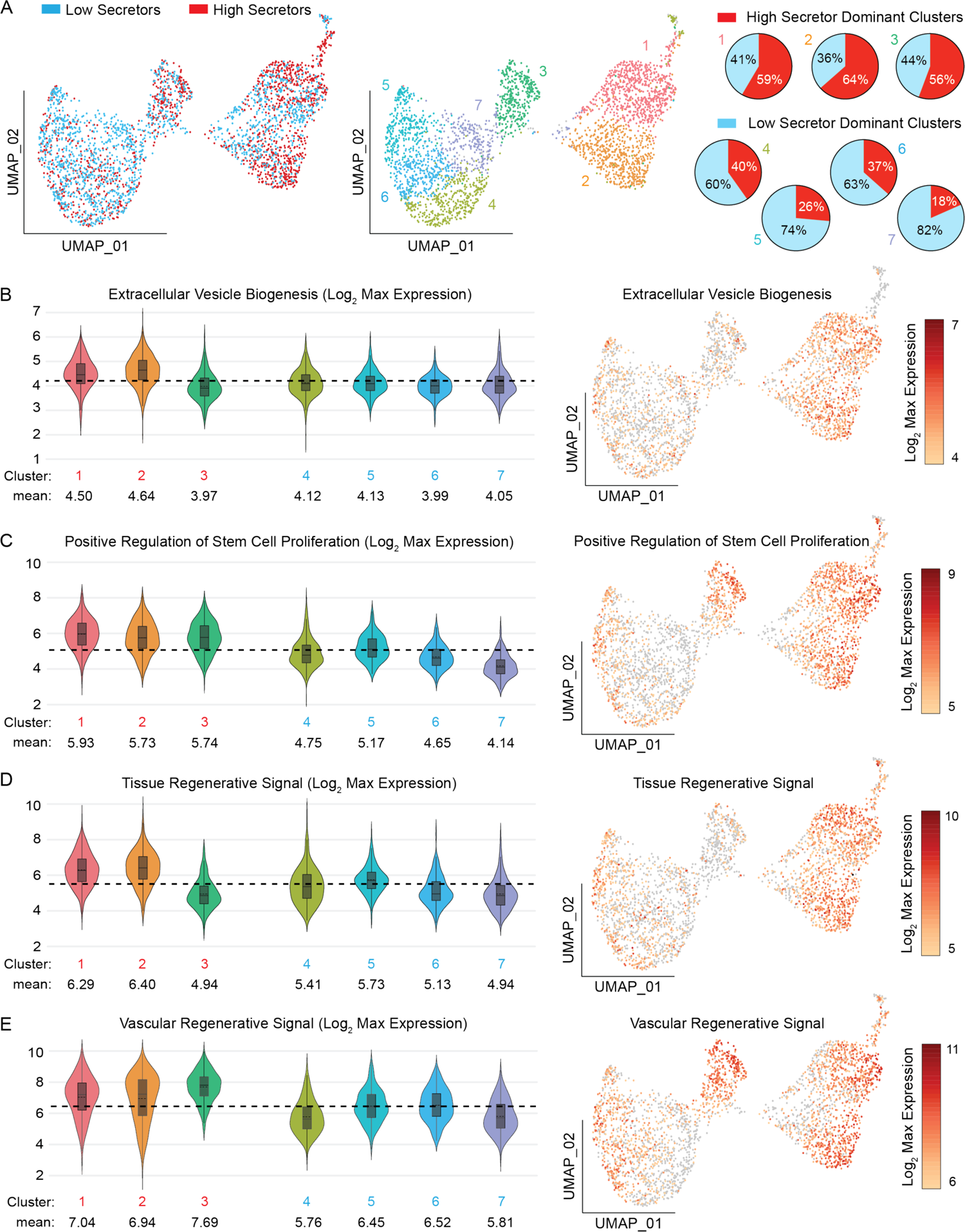
Transcriptomic analysis of low and high EV secreting cells. A) Uniform Manifold Approximation and Projection (UMAP) representation of high (red) and low (blue) secretors. Distribution of high and low secretors in each cluster are represented as pie charts. Clusters 1, 2, 3 are high secretor dominated clusters, while clusters 4, 5, 6, 7 are low secretor dominated clusters. B) Log2 max expression level associated EV biogenesis signature (GO:0140112 Extracellular Vesicle Biogenesis), C) positive regulation of stem cell proliferation (GO:2000648 Positive regulation of stem cell proliferation), and D) tissue regenerative signal or E) vascular regenerative signal represented in each cluster. Black dashed line represents average expression level from all clusters.

To investigate the gene expression profile associated with EV secretion, we first tested whether an EV biogenesis gene ontology signature (GO:0140112 Extracellular Vesicle Biogenesis) marked the clusters with high EV secreting cells specifically. As expected, the EV biogenesis signal was enriched among high secretor dominant clusters (1 and 2) compared to low secretor clusters (Figure 3B). Genes that play a critical role in vesicular trafficking and endosomal recycling processes such as RAS-related GTP binding proteins (RAB7A, RAB27B), tumor susceptibility gene 101 protein (TSG101), hepatocyte growth factor-regulated tyrosine kinase (HGS), and vacuolar protein sorting 4A homolog A (VPS4A) were specifically enriched in these clusters (Supplementary Figure 4B)^10–12^.

As we have observed increased proliferation phenotypes from high EV secretors post-isolation, we further analyzed expression of genes that positively regulate stem cell proliferation (GO:2000648) (Figure 3C). All the high secretor dominant clusters (1, 2, and 3) exhibited increased expression of signatures associated with proliferation in stem cells compared to the low secretor dominant clusters. Some of the upregulated genes in high secretors within this gene ontology were high mobility group A2 (HMGA2), nerve growth factor (NGF), odd-skipped related transcription factor (OSR2), RUNX family transcription factor 2 (RUNX2) and tata binding protein 4 beta (TAF4b) (Supplementary Figure 4C). HMGA2 enhances stem cell proliferation and reduces aging processes by the increased expression of cyclin E and decreased expression of cyclin-dependent kinase inhibitors^13^. NGF promotes MSC proliferation and chondrogenic differentiation through increasing its sensitivity to surrounding stimulating factors^14^.

Since we were interested in applying MSCs in the treatment of cardiac damage, we further analyzed expression of genes that were previously associated with producing EV molecular cargos known for their therapeutic potential to regulate angiogenesis in treating myocardial infarction^15^. Like EV biogenesis signatures, high secretor dominant clusters 1 and 2 exhibited greater expression of these pro-angiogenic factors (Figure 3D). These genes included vascular endothelial growth factor (VEGFA), hepatocyte growth factor (HGF), epidermal growth factor receptor (EGFR), nuclear factor kappa B subunit 1 (NKFB1) and CXC-motif chemokine ligands (CXCL1, CXCL2, CXCL3, CXCL6, CXCL8) (Supplementary Figure 5A). Some of the highly expressed genes are driven by highly expressed transcription factors forming a gene expression node. NFKB1, for example, plays a critical role in driving the expression of other pro-angiogenic genes such as VEGFA and CXCL8^16, 17^. Interestingly, only high EV secretor dominant cluster 3 was specifically enriched with another angiogenesis-inducing component, CXC-motif chemokine ligand 12 (CXCL12), which was not expressed in any other clusters (Supplementary Figure 5). This result demonstrates that the selection of high EV secretors using nanovials can better identify specific functional base cells expected to promote angiogenesis, and ultimately improve the clinical treatment of myocardial infarction.

We recently also identified another regenerative gene expression signature associated with high VEGF-A secretion^18^, expected to also be involved in driving angiogenesis and found the encoding genes for this vascular regenerative signal (VRS) to be more highly expressed in clusters 1, 2, and 3, which were enriched with high secretors. Interestingly, cluster 3, which had reduced EV biogenesis or myocardial infarction regenerative signal mentioned above, showed the highest enrichment in VRS (Figure 3D). Insulin-like growth factor binding protein 6 (IGFBP6) and heme oxygenase 1 (HMOX1) were the most differentially expressed genes in cluster 3 and fibronectin 1 (FN1) was also overexpressed in the high EV secretor dominant clusters (Supplementary Figure 5B). Different gene expression among high EV secreting clusters indicates that there exists transcriptomic heterogeneity even in populations with similar EV secretion levels, suggesting that gene expression of surface marker proteins alone may not be sufficient to mark high EV secretor populations.

Among the top 20 overexpressed transcripts in high secretors, ankyrin repeat and sterile alpha motif domain (ANKS1B), Tenascin XB (TNXB) and platelet endothelial cell adhesion molecule-1 (PECAM1) were the most notable due to their involvement in cell-to-cell signaling, vesicle transportation, extracellular matrix organization and cell migration (Supplementary Figure 6A)^19–21^. The top 20 up-regulated genes in high secretors were also associated with biological processes related to tissue development, cell migration and regulation of signaling receptor activity, such as endothelial cell development (GO:0001885), cell-cell adhesion (GO:0098609) and neurotransmitter receptor activity regulation (GO:0099601) (Supplementary Figure 6B).

### High secretors increase the viability of injured cardiomyocytes via EV-mediated paracrine repair

To evaluate whether selection of cells based on EV secretion would lead to differences in MSC-mediated tissue repair, we sorted mouse MSCs based on EV secretion levels using nanovials (Supplementary Figure 7). Cell proliferation and EV secretion were both increased in high-secreting (high-sec) mouse cells compared to low-secreting (low-sec) cells as was found for human cells. To compare the paracrine repair ability of these MSCs, we isolated EVs from 1 million cells of each type and incubated them with H9C2 cardiac cells after exposure to hydrogen peroxide. EVs from high-sec MSCs were able to better rescue cells from oxidative stress mediated apoptosis (TUNEL positive) when compared to the EVs isolated from the same number of low-sec MSCs, which implied an increased paracrine repair activity of high-sec MSCs on a per cell basis (Supplementary Figure 8).

### High EV secretors have greater therapeutic potential in a mouse myocardial infarction model

To test whether the augmented paracrine activity would translate to increased therapeutic potency *in vivo*, we performed a head-to-head comparison of the high-sec MSCs to the low-sec MSCs in cardiac repair using a mouse myocardial infarction (MI) model (Figure 4A). Before injecting cells into the injured heart, mouse MSCs were suspended in 10 mg/mL hyaluronic acid (HA) hydrogel, acting as a delivery medium to aid cell retention in the heart after injection. As our previous study indicated that intra-pericardial cavity (IPC) injection achieved a higher stem cell engraftment and resulted in a boosted paracrine activity mediated by the stem-cell secreted EVs^22–24^, we injected high-sec and low-sec MSCs using IPC injection and the same amount of empty HA hydrogel was injected as the control. We monitored changes in the cardiac function one day before, 2 days, 14 days and 28 days after the surgery (Figure 4A) by calculating left ventricular ejection fraction (LVEF) and left ventricular fractional shortening (LVFS) from echocardiography (Figure 4B). A sharp decrease in LVEF and LVFS at 2 days indicated successful MI in all groups. At Day 14, both LVEF and LVFS started to recover in both high-sec MSC and low-sec MSC treatment groups but not in the control group (Figure 4C and D). At Day 28, we observed significant differences in the overall left ventricular functions of all experimental groups, with the high-sec MSC treatment group having the highest LVEF and LVFS (Figure 4C and D). In addition, the cardiac functions of the low-sec MSC treatment group showed a continuous increase throughout the observation period but failed to recover to the baseline level. To further examine the linkage between the EV-secreting ability and therapeutic potential, we referred to our previous study where wild-type MSCs were injected using the same delivery method^22^. Interestingly, the results showed that the 28-day LVEF and LVFS in the wild-type MSC treatment group were intermediate between those of the high- and low-sec MSC treatment groups. This suggests that the therapeutic potential of wild-type MSCs is lower than that of high-sec MSCs, but higher than that of low-sec MSCs, presumably due to the difference in EV-mediated paracrine activity.

**Figure 4.**
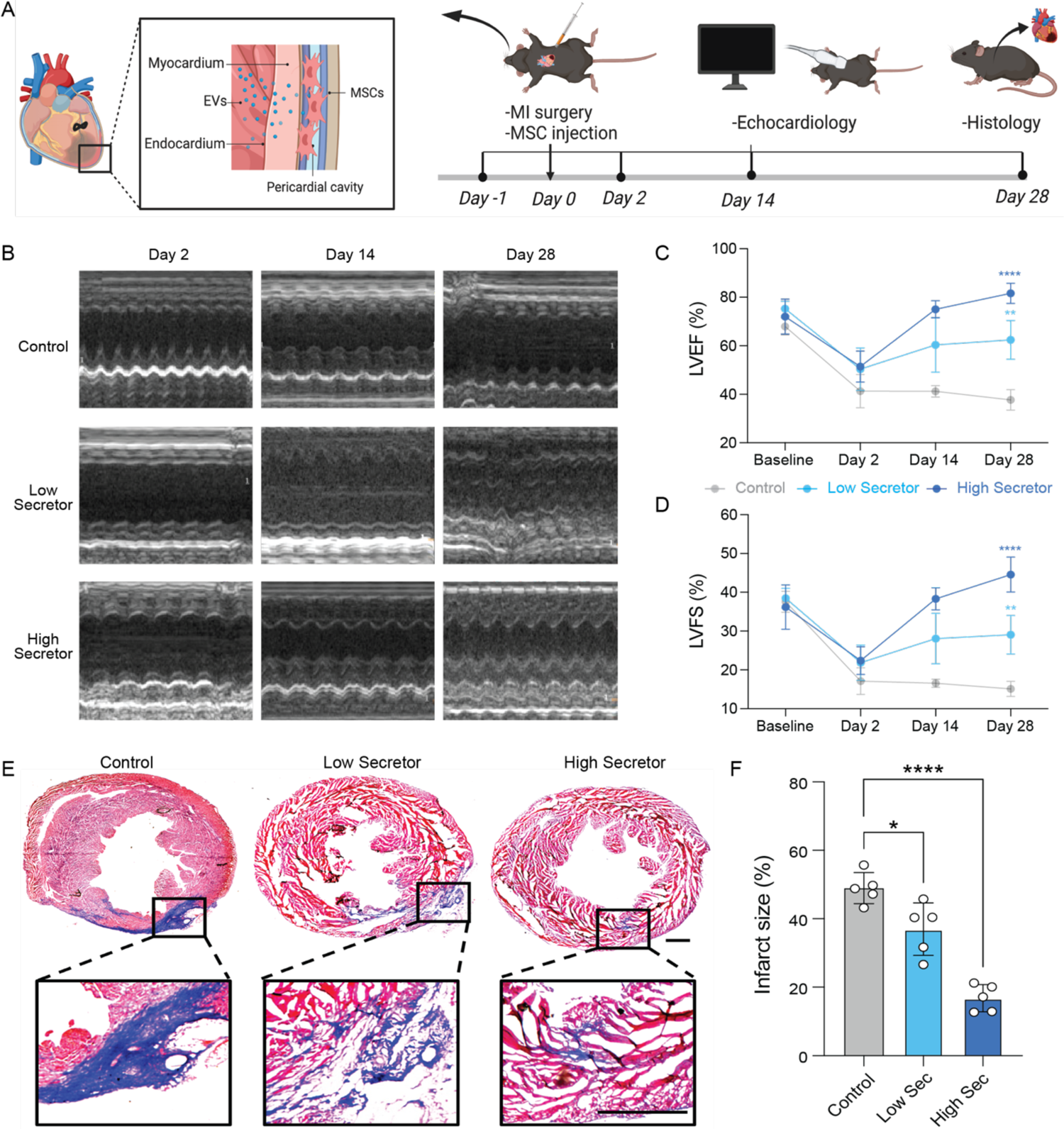
High EV secreting (high-sec) MSC therapeutic potential compared to low EV secreting (low-sec) MSCs in a myocardial infarction (MI) model. A) Schematic illustration of the *in vivo* study design. MSCs were injected into the intra-pericardial cavity (IPC) after MI surgery. Echocardiography was performed at 2 days, 14 days and 28 days after the surgery. B) Representative M-mode echocardiography images at 2 days, 14 days and 28 days post MI surgery. C) LVEF of mice receiving different treatments at 2 days, 14 days and 28 days post MI surgery (n = 5, biological replicates). D) LVFS of mice receiving different treatments at 2 days, 14 days and 28 days post MI surgery (n = 5, biological replicates). E) Representative Masson’s trichrome staining of heart sections (red = healthy tissue; blue = scar) 4 weeks after treatments. Scale bar: 1 mm. F) Quantified infarct size of heart sections. All data are means ± SD. Comparisons between more than two groups were performed using the one-way analysis of variance (ANOVA), followed by Tukey’s honestly significant difference (HSD) post hoc test. The comparisons between samples are indicated by lines, and the statistical significance is indicated by asterisks above the lines.

The histology analysis of heart sections provided further evidence for the enhanced cardiac repair capability of high-sec MSCs. Specifically, the high-sec MSC treatment group exhibited the smallest infarct size, which was significantly lower than that of both the low-sec MSC treatment group and the control group (Figure 4E and F). In addition, high-sec MSCs were found to reduce cell apoptosis induced by MI and promote cell proliferation in the heart. This was evidenced by a decrease in caspase-3 expression and an increase in the number of Ki67-positive cells (Figure 5A and B). Furthermore, an increased density of CD31-positive blood vessels was found in the heart after high-sec MSC treatment (Figure 5C). Altogether, high-sec MSCs can better mitigate MI injury and help to restore left ventricular function by reducing cell death, promoting cell proliferation, and increasing vascular density around the infarct region.

**Figure 5.**
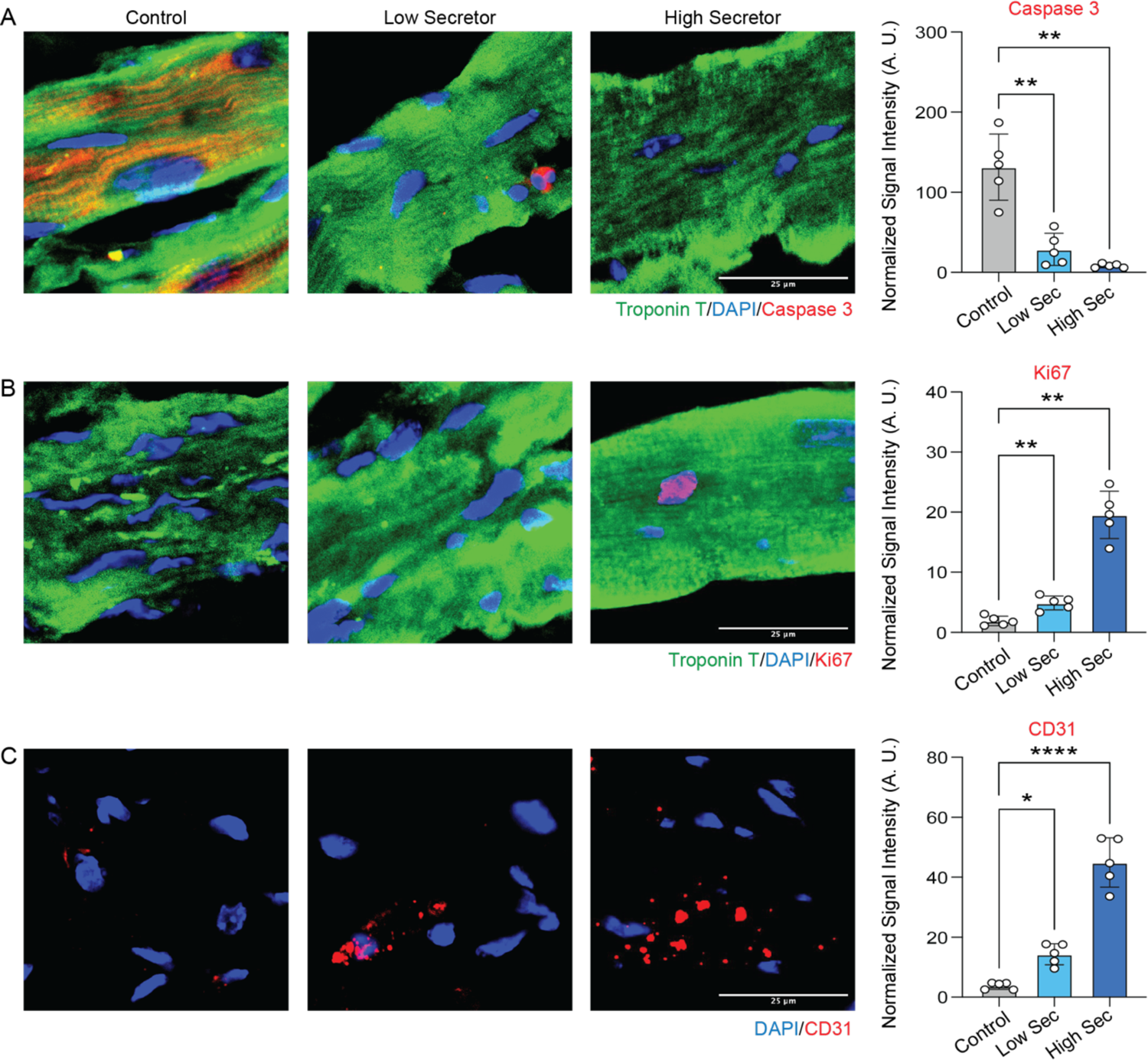
IHC staining of heart sections. A) Representative fluorescence images of cell apoptosis determined by caspase-3 expression (red). Scale bars, 50 μm. Quantitation of normalized caspase-3 levels in cells (n = 5, biological replicates). B) Representative fluorescence images showing Ki67+ cells (red) in the myocardium. Scale bars, 50 μm. Quantitation of normalized Ki67 expression in cells (n = 5, biological replicates). C) Representative fluorescence images of vascular regeneration indicated by CD31 staining (red). Scale bars, 100 μm. Quantitation of normalized CD31 staining in cells (n = 5, biological replicates). All data are means ± SD. Comparisons between more than two groups were performed using the one-way analysis of variance (ANOVA), followed by Tukey’s honestly significant difference (HSD) post hoc test. The comparisons between samples are indicated by lines, and the statistical significance is indicated by asterisks above the lines.

### High EV secreting MSCs treatment exhibits excellent safety in vivo

Next, we assessed the safety of using high-sec and low-sec MSCs for treating MI. At the experimental endpoint, we harvested major organs (lung, liver, spleen and kidney) from mice and conducted histology evaluation using hematoxylin and eosin (H&E) staining (Supplementary Figure 9A). There were no significant differences in the lungs, livers, spleens, and kidneys of the treated mice compared to the healthy mice, suggesting that high-sec MSC injection does not cause severe side effects such as tumor growth. The blood chemistry results also confirmed that the liver and kidney functions were not affected by high-sec MSC treatment (Supplementary Figure 9B). Interestingly, we observed that the high-sec MSC treatment group had lower levels of ALT and AST compared to the low-sec MSC treatment group, which could be due to the increased paracrine activity of high-sec MSCs that helps improve liver functions.

## Discussion and Conclusion

Although there has been increasing interest in using cells that produce EVs therapeutically, our work is the first to explore how selecting sub-populations of cells that secrete more EVs affects therapeutic outcomes. We found that delivering enriched populations of cells that secrete more EVs leads to improved *in vitro* protection and *in vivo* functional and structural recovery after a myocardial injury. These results directly link the EV-secretory capacity of cells to therapeutic outcome and provide further evidence for the role of increased EV quantity in recovery from injury.

Notably, in selecting sub-populations of high EV secretors, we also found that this phenotype was maintained and appeared heritable across multiple cell doublings, indicating a conserved epigenetic program present in a fraction of MSCs that appears to define a more regenerative sub-type of cells that also proliferates more rapidly. This phenotype is reflected in single-cell transcriptomic measurements which show increased expression of genes previously linked to regeneration and vascular growth, including the vascular regenerative signal (VRS). VRS marks a population of cells that also have high secretion of other factors important for tissue regeneration, like vascular endothelial growth factor A (VEGF-A)^25^. Given this data, there is a possibility that high EV secretion serves as a marker for a more regenerative and secretory phenotype of cells, and EV quantity itself may not be solely responsible for the therapeutic benefit we observed.

In order to select sub-populations of EV-secreting cells, we needed to develop an assay to measure and sort cells based on EV release at the single-cell level. Multiple studies recently reported immunoaffinity microwell-based platforms for analyzing EV secretion at the single-cell level, but currently no technology enables the retrieval of cells^26–28^. Sorting viable intact cells is important for further downstream regrowth or analysis (e.g. single-cell RNA sequencing). Using nanovials we were able to select single-cells based on EV secretion level using commercially available instruments and reagents (Sony SH800S 130 micron chip) and directly process cells for transcriptomic analysis using the 10X Genomics single-cell RNA sequencing platform to uncover factors associated with the high EV secretion phenotype. Further studies can quickly build on our results leveraging instruments and nanovials (Partillion Bioscience) that are all available commercially.

Our approach to measure EVs focused on EVs marked with two tetraspanins (CD63 and CD9), however, by adjusting the capture and detection antibodies, different or multiple types of EVs can be captured and analyzed on nanovials. For example, multiple detection antibodies could be applied with different fluorophores and/or the nanovial surface could be conjugated with more than one capture antibody targeting different EV-surface presented proteins, including engineered targeting protein^29^. We have previously shown the ability to multiplex the detection of secreted cytokines^30^. By further characterizing the types of EVs released, researchers could further sub-categorize cells based on the propensity to secrete subsets of EVs and refine our understanding of what are the most important sets of EVs to be released for therapeutic benefit.

The ability to sort cells secreting therapeutically active substances directly can have a large impact on the manufacturing and quality control of EV and cell therapies. Out of 33 registered EV-based clinical trials, 20 clinical trials utilize MSC-derived EVs, where three trials specifically aimed to transplant engineered MSCs for overproduction of EVs^31^. Using nanovials, we can select the most potent cells for growth and delivery, alleviating the challenges associated with significant EV secretion heterogeneity, and batch to batch variations in current therapeutics. Given that EV secretion is sustained across multiple cell doublings, we can also select clones for master and working cell banks based on increased EV productivity for direct EV-based therapeutics. Finally, characterizing EV secretion amount and heterogeneity of a finished cell therapy product (e.g. using ImageStream imaging cytometry) may provide a more direct potency assay that ties cell function to therapeutic efficacy, improving the reproducibility and patient outcomes for each therapeutic batch.

## Materials and Methods

### Nanovial fabrication

Polyethylene glycol biotinylated nanovials with 50 μm diameters were fabricated using a three-inlet flow-focusing microfluidic droplet generator, sterilized and stored at 4°C in Washing Buffer consisting of Dulbecco’s Phosphate Buffered Saline (Thermo Fisher) with 0.05% Pluronic F-127 (Sigma), 1% 1X antibiotic-antimycotic (Thermo Fisher), and 0.5% bovine serum albumin (Sigma) as previously reported ^32^.

### Nanovial Functionalization

#### Streptavidin conjugation to the biotinylated cavity of nanovials

50 μm sterile nanovials were diluted in Washing Buffer five times the volume of the nanovials (i.e. 100 μL of nanovial volume was resuspended in 400 μL of Washing Buffer). A diluted nanovial suspension was incubated with equal volume of 200 μg/mL of streptavidin (Thermo Fisher) for 30 minutes at room temperature on a tube rotator. Excess streptavidin was washed out three times by pelleting nanovials at 2000xg for 30 seconds on a Galaxy MiniStar centrifuge (VWR), removing supernatant and adding 1 mL of fresh Washing Buffer.

#### Anti-CD63 EV capture antibody labeled nanovials

50 μm streptavidin-coated nanovials were reconstituted at a five-time dilution in Washing Buffer containing 270 nM (40 μg/mL) of biotinylated anti-CD63 antibody (Biolegend, 353018). Nanovials were incubated with antibodies for 30 minutes at room temperature on a rotator and washed three times as described above. Nanovials were resuspended at a five times dilution in Washing Buffer or culture medium prior to each experiment.

### Cell culture

#### Immortalized mesenchymal stem cells (iMSC)

hTERT immortalized adipose derived MSCs (ATCC, ASC52telo) were cultured based on manufacture’s protocol using MSC basal media (ATCC, PCS500030) and growth kit (PCS500040). For every secretion assay in this study, cells at passage 18 were used to maintain consistency.

C57BL/6 Mouse Bone Marrow Mesenchymal Stem Cells. Mouse mesenchymal stem cells (cellbiologics, C57-6043) were maintained in Iscove’s modified Dulbecco’s medium (IMDM) (Invitrogen) containing 10% fetal bovine serum (FBS). Cells at passage 8 were used in the *in vivo* study.

H9C2 cells (ATCC, CRL-1446) were maintained in Iscove’s modified Dulbecco’s medium (IMDM) (Invitrogen) containing 10% fetal bovine serum (FBS). Cells at passage 6 were used in the *in vitro* study.

### Nanovial secretion assay general procedure

#### Cell loading onto nanovials

Each well of a 24-well plate was filled with 1 mL of media and 50 μL of 5X reconstituted functionalized nanovials (10 μL of nanovial volume=100,000 total nanovials) was added in each well using a standard micropipette. Cells were seeded in each well and extra culture medium was added to make a total volume of 1.5 mL. Each well was mixed by simply pipetting 5 times with a 1000 μL pipette set to 1000 μL. The well plate was transferred to an incubator to allow cell binding; the volume in each well was pipetted up and down again 5 times with a 200 μL pipette set to 200 μL at 30-minute intervals. After one hour, nanovials were strained using a 20 μm cell strainer to remove any unbound cells and recovered. During this step, any unbound cells were washed through the strainer and only the nanovials (with or without cells loaded) were recovered into a 12-well plate with 2 mL of media by inverting the strainer and flushing with media.

#### Secretion accumulation and secondary antibody staining on nanovials

After cell loading, cells on nanovials were incubated for 24 hours in the incubator to accumulate secretion. Each sample was recovered in a conical tube with 5 mL wash buffer and centrifuged for 5 minutes at 200xg. Supernatant was removed and nanovials were reconstituted at a ten-fold dilution in Washing Buffer containing detection antibody and/or cell viability dye (calcein AM) to label secreted EVs and viability of cells. For each 100,000 nanovials (∼ 10 μL nanovial volume), 5 μL of anti-CD9 BV650 antibody with 85 μL of 0.3 μM Calcein AM solution, unless otherwise stated. Nanovials were incubated with the detection antibody at 37°C for 30 minutes, protected from light. After washing nanovials with 5 mL of Washing Buffer, nanovials were resuspended at a 40-fold dilution in Washing Buffer and transferred to a flow tube.

#### Flow cytometer analysis and sorting

All flow cytometry sorting were performed using the SONY SH800 cell sorter equipped with a 130 micron sorting chip (SONY Biotechnology). The cytometer was configured with violet (405 nm), blue (488 nm), green (561 nm) and red (640 nm) lasers with 450/50 nm, 525/50 nm, 600/60 nm, 665/30 nm, 720/60 nm and 785/60 nm filters. Standard gain settings for different sensors are indicated in table 2 below and gains were adjusted depending on the fluorophores used. In each analysis, samples were compensated using negative (blank nanovials) and positive controls (purified EV conjugated nanovials labeled with each fluorescent detection antibody or cells stained with calcein AM). Nanovial samples were diluted to approximately 623 nanovial/μL in Washing Buffer for analysis and sorting. Drop delay was configured using standard calibration workflows and single-cell sorting mode was used for all sorting as was previously determined to achieve the highest purity and recovery ^33^. A sample pressure of 4 was targeted. The following order of gating strategy was used to identify cells on nanovials with strong secretion signal: 1) nanovial population based on high forward scatter height and side scatter area, 2) calcein AM positive population, 3) CD9 EV secretion signal positive population based on fluorescence peak area and height. For analyzing sample using ImageStream imaging-based flow cytometer, 10 μL nanovial sample was resuspended in 200 μL Washing buffer and analyzed based on manufacture’s protocol. Images were analyzed by IDEAS software distributed by Luminex.

### EV secretion phenotype analysis using ExoView

Cells were seeded in T-75 flask at seeding density of 0.2 million cells per flask with stem cell basal media with 2% FBS. After 24 hours of initial seeding, media was changed to OPTI-MEM I reduced serum media without phenol red (Fisher Scientific, 11-058-021). After 48 hours, conditioned media was centrifuged at 3000 RPM for 10 minutes at 4°C and the supernatant was filtered using 0.2 μm syringe filter (VWR, 28143-310). Sample placed in ultracentrifuge tubes (Fisher Scientific, NC9732446) was placed into pre-cooled 50 Ti fixed angle titanium rotor and centrifuged in Optima XPN-100 Ultracentrifuge at 40,000 RPM for 120 minutes at 4°C. Media was removed and the pellet was resuspended in 100 μL cold PBS. Sample was analyzed by ExoView R100 for CD63, CD9 and CD81 markers based on the manufacture’s protocol.

### Dynamic range of CD63+CD9+ EVs on nanovials

Nanovials were labeled with biotinylated anti-CD63 antibody using the modification steps mentioned above and incubated with EVs isolated from ultracentrifugation at 0, 30, 60, or 120 μg/mL for 2 hours at 37°C. Excess EVs were removed by washing nanovials three times with Washing Buffer. Nanovials were pelleted at the last wash step and incubated with anti-CD9 as described in secondary antibody staining procedure above. Following washing three times, nanovials were reconstituted at a 50 times dilution in the Washing Buffer and transferred to a flow tube. Fluorescent signal on nanovials was analyzed using a cell sorter with sensors and imaged using fluorescence microscope.

### Single-MSC loading and statistics

50 μm nanovials labeled with anti-CD63 antibodies were prepared using the procedures described above. To test cell concentration dependent loading of nanovials, 0.1 x 10^6^ (1 cell per nanovial) and 0.2 x 10^6^ (2 cells per nanovial) of cell tracker deep green stained cells were each seeded onto 100,000 nanovials in a 24-well plate and recovered as described above. Samples resuspended at 40-fold dilution in Washing buffer were analyzed using SONY sorter for loading efficiency. The population in each gate (single-cell loaded nanovial, multiple-cell loaded nanovial, nanovial aggregate) was sorted into 96-well plate containing Washing buffer and imaged using fluorescence microscope.

### Analysis and sorting of single-cells based on CD63+CD9+ EV secretion level

0.1 million MSCs were loaded onto anti-CD63 labeled nanovials and secretion was accumulated for 24 hours. Same number of cells were also loaded onto unlabeled nanovials (no capture antibody) as a negative control. After 24 hours secreted EVs were labeled with fluorescent anti-CD9 antibodies and calcein AM viability dye. After resuspending nanovials at 40-fold dilution in Washing Buffer, a small fraction of sample was transferred to a 96-well plate to be imaged using a fluorescence microscope prior to sorting. Samples were analyzed using a cell sorter based on a combination of fluorescence area and height signals. To sort live single cells based on secretion signal, nanovials with calcein AM staining were first gated and high, medium, or low secretors and sorted by thresholding above the negative control sample on CD9 fluorescence area and height signals. Sorted samples were imaged with a fluorescence microscope to validate the enrichment of nanovials based on the amount of secreted EVs captured on the nanovials.

### Expansion of cells sorted based on EV secretion level

In each well of a 96-well plate filled with 150 μL stem cell basal media, a single-cell loaded nanovial was sorted based on EV secretion level as low, medium and high EV secretors. 3000 cell-loaded nanovials were also sorted as bulk-sorted population in each well of a 96-well plate. Since low secretors proliferated much slower than medium and high secretors, cells were expanded till Day 13 and re-seeded into T-75 flask at 0.2 million cells per flask. Similarly, single-cell colony was expanded till Day 22 and re-seeded into T-75 flask at the same seeding density. 24 hours after cells are seeded, media was changed to OPTI MEM reduced serum media. After 48 hours (Day 25 for single-cell colony and Day 16 for bulk-sorted population), conditioned media was collected, and the final number of cells were counted. EVs were collected from conditioned media via ultracentrifugation and quantified by ZetaView analysis at University of North Carolina Nanomedicines Characterization Core Facility. For calculating EV production efficiency, the total EV count was divided by the total number of cells in each sample.

### Single-cell transcriptomic analysis

The standard protocol for 10X Chromium single cell 3’ GEX was followed unless otherwise noted. Sorted samples reconstituted at 18 μL were loaded into the 10X Chromium Next GEM Chip for partitioning each nanovial or cell into droplets allowing the PCR amplification and enrichment of gene expression for individual cell barcoded cDNA. Single-cell libraries were constructed using the manufacturer-recommended protocol by the UCLA Technology Center for Genomics & Bioinformatics. Libraries were then sequenced on Novaseq SP (235-400M/lane) (Illumina). The Cell Ranger pipeline was used for sample de-multiplexing. 10X Loupe browser was used for further analysis and generation of UMAP and violin plots. For EV biogenesis (GO: 0140112) and stem cell proliferation signatures (GO: 2000648), gene list was downloaded from the Jackson Laboratory. For tissue regeneration and vascular regenerative signatures, gene list was retrieved from previously studies ^15, 18^.

#### *In vitro* ischemic injury model

To establish the ischemic injury *in vitro*, H9C2 cells were incubated with 500 μM H2O2 in complete medium for 2 hours. After that, the H2O2 containing medium was replace by complete medium with exosomes isolated from same number of MSCs with different secretion abilities. For 24 hour treatment, the cell apoptosis was evaluated using TUNEL staining^34^.

### IPC injection of MSCs in mice model of MI

The Institutional Animal Care and Use Committee (IACUC) and NIH Guide for the Care and Use of Laboratory Animals were adhered to in all animal research. To create a mouse MI model, mice were anesthetized through intraperitoneal injection of K-X cocktail (100 mg/kg ketamine and 10 mg/kg xylazine), while a small animal ventilator (SAR-1000 Small Animal Ventilator, CWE, Inc.) provided artificial ventilation as life support. The LAD coronary artery was ligated permanently under sterile conditions to induce ischemia. Following MI induction, 0.2 million cells with 10 µL 10 mg/mL hydraulic acid (HA) hydrogel were injected into the pericardial cavity of each mouse heart. The chest was then closed, and the animal was permitted to recover.

### Mouse echocardiology

Echocardiography was carried out at various time points, including baseline (prior to MI surgery), and 2, 14, and 28 days after the MI procedure. To conduct transthoracic echocardiography, mice were anesthetized using an isoflurane/oxygen mixture and positioned supine. A 40-MHz probe from the high-frequency ultrasound system (Prospect, S-Sharp, New Taipei City, Taiwan) was used to obtain B (brightness, 2D) and M (motion) mode images. The images were recorded and assessed blindly. The ejection fraction was calculated using the following formula: EF = (LVEDV - LVESV/LVEDV) × 100%, while the fractional shortening was determined using FS = (LVEDD - LVESD/LVEDD) × 100%.

### Histology

Mice tissues were harvested and fixed using a neutral buffered 10% formalin solution (NBF, Sigma-Aldrich, St. Louis, MO, USA) overnight. Afterward, the tissues were dehydrated in a 30% sucrose solution at 4°C for at least one night. Subsequently, the tissues were cryopreserved in optimal cutting temperature (OCT) compound and cryosectioned (CryoStats, Leica). Pathological assessments were carried out using H&E and Masson’s trichrome staining in accordance with the manufacturer’s instructions (Sigma-Aldrich, St. Louis, MO, USA).

For immunofluorescence staining, cryosections of the tissues were first permeabilized and blocked using a protein block solution (Dako, Carpinteria, CA, USA) containing 0.1% saponin (Sigma-Aldrich, St. Louis, MO, USA). The primary antibodies were then added and incubated overnight at 4°C, including rabbit anti-Caspase 3 (1:100; ab184787, Abcam, Cambridge, UK), rabbit anti-Ki67 (1:100; ab15580, Abcam), anti-CD31 antibody (1:100; EPR17259, Abcam), and mouse Anti-Cardiac Troponin T antibody (1:100; ab8295, Abcam) to target proteins of interest. Fluorophore-conjugated secondary antibodies were then added for fluorescent imaging. All the tissue slides were mounted by ProLong gold antifade mountant with DAPI (Thermo Fisher, US) before being imaged using an epifluorescent microscope from Olympus.

### Serum chemistry analysis

At the experimental endpoint, blood samples were collected from mice, and serum was isolated using BD Vacutainer® serum collection tubes (BD, 367814). The serum chemistry test was performed by the Department of Clinical Pathology, College of Veterinary Medicine, NC State University.

## Acknowledgements

We would like to thank other members of Di Carlo lab and Ke Cheng lab for helpful comments and discussion in preparation of this manuscript. We thank UCLA Jonsson Comprehensive Cancer Center (JCCC) and Center for AIDS Research Flow Cytometry Core Facility. We thank UCLA Technology Center for Genomics & Bioinformatics for performing sequencing services. We also thank Robert Knight and Sophie Deng for allowing us to use ExoView.

## Author contributions

Conceptualization: DK, DD, XC, KC Methodology: DK, DD, JK, XC, KC Investigation: DK, XC, SU, DZ, JL, NT, SH Formal Analysis: DK, XC, BH Visualization: DK, XC

Funding acquisition: DK, DD

Writing – original draft: DK, XC, DD, KC

Writing – review & editing: DK, XC, DD, KC, JK, SU, DZ, JL

