## Supplementary Information for "Optimizing Cell Therapy by Sorting Cells with High Extracellular Vesicle Secretion"

**Affiliations:**
[1]Department of Bioengineering, University of California, Los Angeles; Los Angeles, CA 90095, USA.
[2]Joint Department of Biomedical Engineering, University of North Carolina at Chapel Hill and North Carolina State University; Chapel Hill, NC 27599, and Raleigh, NC 27607, USA.
[3]Department of Molecular Biomedical Sciences and Comparative Medicine Institute, North Carolina State University; Raleigh, NC 27607, USA.
[4]Cytek Biosciences; Fremont, CA 94538, USA.
[5]Department of Pathology and Laboratory Medicine, University of Pennsylvania; Philadelphia, PA 19104, USA.
[6]Department of Bioengineering, University of Pennsylvania; Philadelphia, PA 19104, USA.
[7]Jonsson Comprehensive Cancer Center, University of California, Los Angeles; Los Angeles, CA 90095, USA.
[8]Department of Mechanical and Aerospace Engineering, University of California, Los Angeles; Los Angeles, CA 90095, USA.
[9]California NanoSystems Institute; Los Angeles, CA 90095, USA.
*These authors contributed equally to this work.

**Supplementary Materials List:**

Supplementary Figure 1. Nanovial fabrication and functionalization.

Supplementary Figure 2. Optimization of single-cell EV secretion assay using nanovials.

Supplementary Figure 3. Regrowth of cells following sorting based on EV secretion levels.

Supplementary Figure 4. Transcriptomic analysis of MSC markers, EV biogenesis and stem cell proliferation expression.

Supplementary Figure 5. Differences in tissue regeneration signature expression between high and low secretors.

Supplementary Figure 6. Gene ontology annotations for top 20 upregulated genes in high secretors.

Supplementary Figure 7. Analysis and isolation of mouse MSCs based on EV secretion level.

Supplementary Figure 8. High secretors exhibit higher potential to reduce cell apoptosis following H2O2-induced rat cardiomyocyte (H9c2 cells) injury.

Supplementary Figure 9. Safety of high-sec and low-sec MSC treatment.

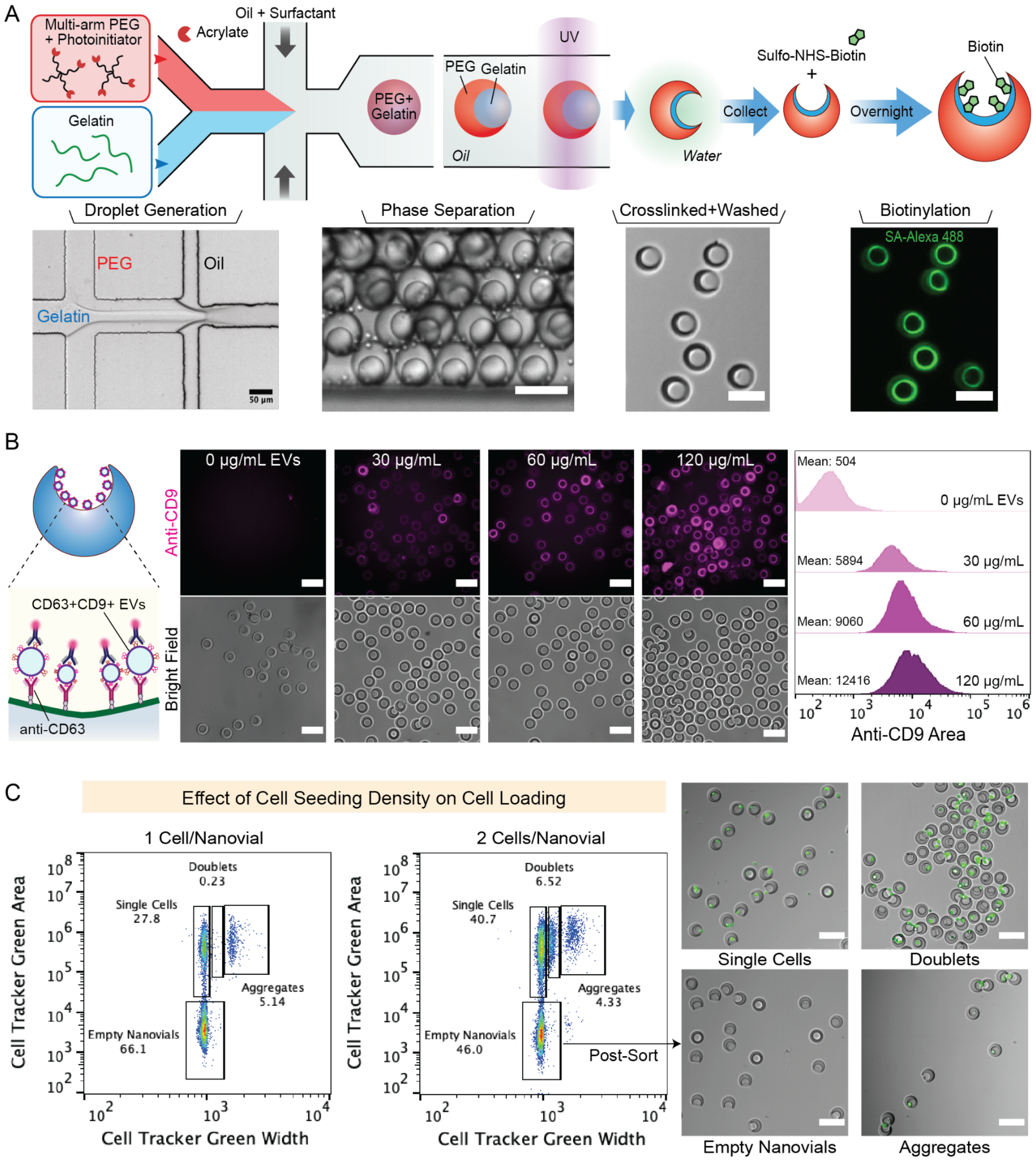


Supplementary Figure 1. Nanovial fabrication and functionalization. A) An aqueous phase consisting of 4-arm-polyethylene glycol (PEG) acrylate and photo-initiator is co-flowed with a gelatin solution in a microfluidic droplet generator. After droplet formation, PEG and gelatin undergo phase separation forming a PEG-rich surrounding phase and gelatin-rich internal phase. The PEG outer structure is cross-linked by exposure to UV light. The gelatin phase is washed away except for a layer that is cross-linked at the interface of the cavity. After collection, nanovials are incubated with sulfo-NHS-biotin to biotinylate the internal gelatin-coated cavity. Localized fluorescence of AlexaFluor488-labeled streptavidin is observed in biotinylated nanovial cavities. Scale bar represents 50 µm. B) Schematic of the EV capture assay shows the anti-CD63 capture and fluorescent anti-CD9 detector antibodies. Flow cytometry fluorescence histograms and fluorescence microscopy images of nanovials following exposure to increasing amounts of EVs. Nanovials were functionalized with anti-CD63 antibodies and incubated with 0, 30, 60, or 120 μg/mL of isolated EVs from the conditioned medium. C) Gating strategy for flow cytometry analysis and sorting of single cells on nanovials. Flow cytometry scatter plots, gates, and microscopy images of gated events are shown. The highest fraction of single-cell loaded nanovials was achieved when cells were seeded at 1 cell per nanovial. Nanovials with single cells were sorted based on the fluorescent area vs. width signal of cells (cell tracker green). Scale bar represents 100 μm.

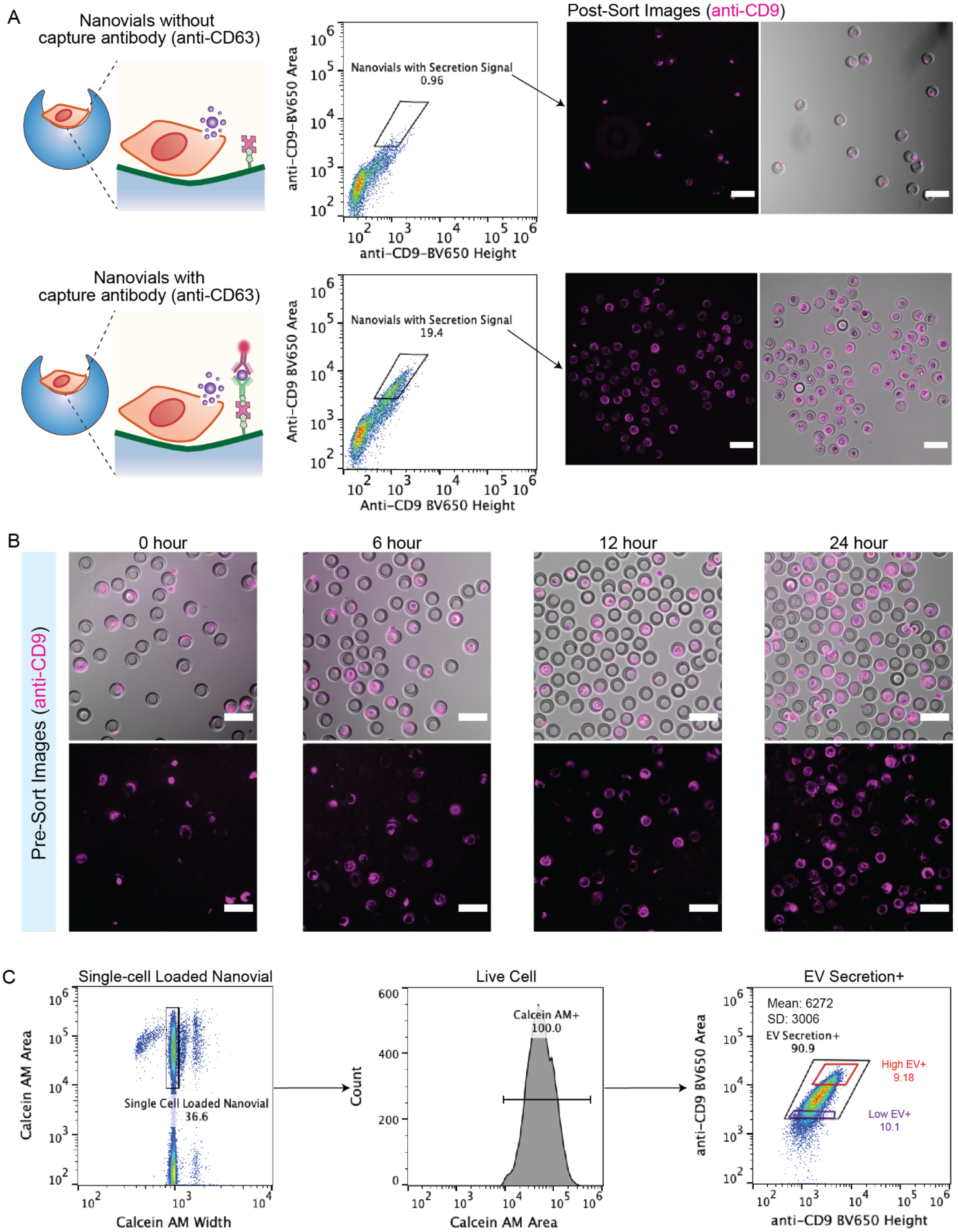


Supplementary Figure 2. Optimization of the single-cell EV secretion assay using nanovials. A) Flow cytometry analysis of EV secretion after 24 hours of secretion accumulation on cell-loaded nanovials. Strong CD9 signal was observed from cells loaded onto nanovials with anti-CD63 EV capture antibody, while secretion was not detected from cells loaded onto nanovials without capture antibody. Fluorescence microscopy images showing post-sort population from the illustrated “secretion positive” gate are shown. Scale bars represent 100 μm. B) Fluorescence microscopy images of EV secretion signals on cell-loaded nanovials after 0 to 24 hours. The highest CD63+CD9+ EV secretion signal was observed when secretion was accumulated over 24 hours on nanovials. Scale bars represent 100 μm. C) Flow cytometry flows showing gating of single-cell loaded nanovials with positive calcein AM staining For viable cells, significant heterogeneity in secretion is observed. Single live cells on nanovials spanned secretion signals over one order of magnitude with coefficient variance of 0.49.

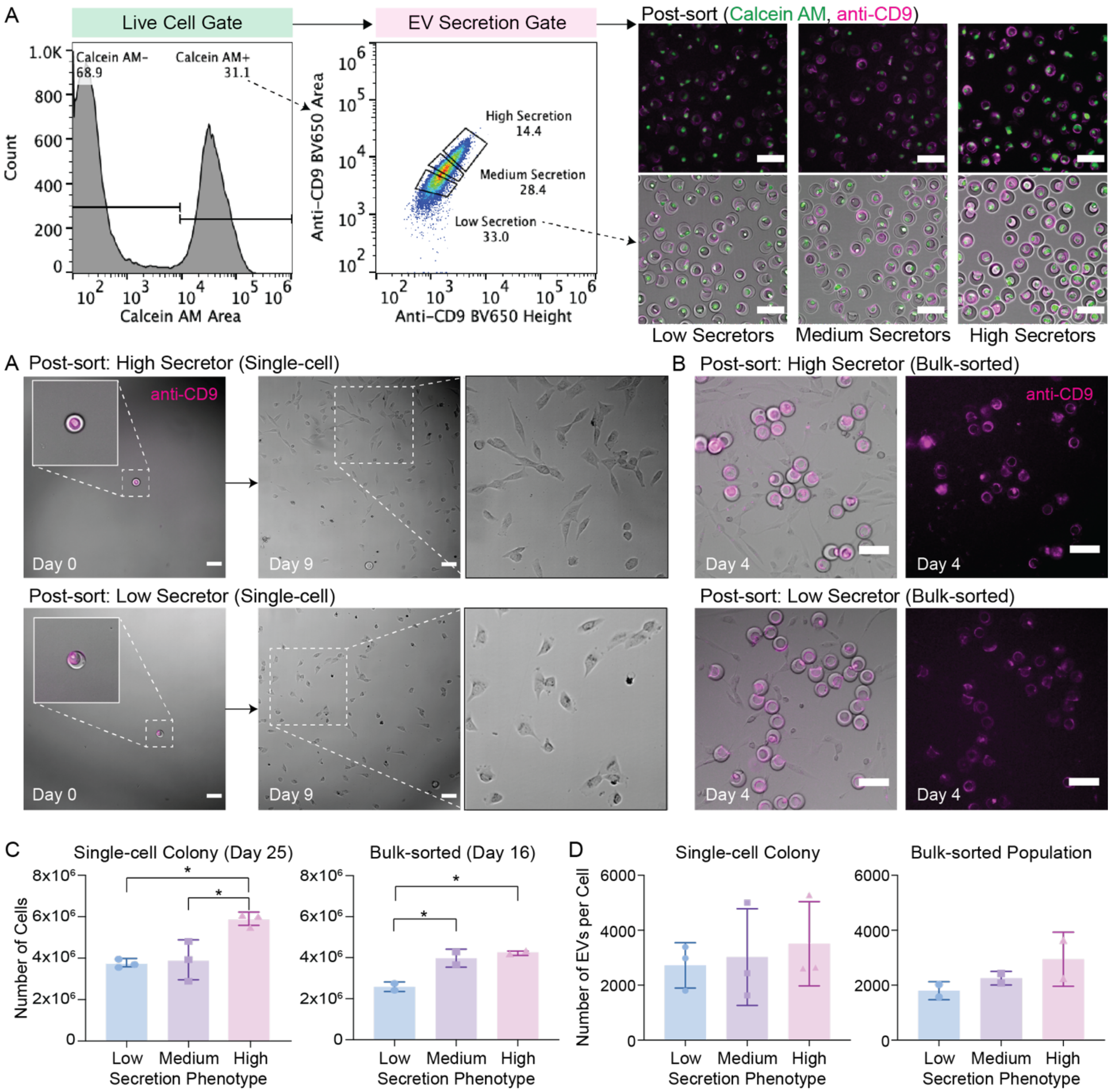


Supplementary Figure 3. Regrowth of cells following sorting based on EV secretion levels. A) Expansion of single-cell colonies from high and low EV secretors shown from Day 0 to Day 9. Scale bars represent 100 µm. B) Expansion of bulk-sorted cells (3000 cells) sorted based on secretion level on Day 4. Secreted EVs are still retained inside the cavity of nanovials at Day 4. C) Final number of cells in a colony expanded from single cells on Day 25 or bulk-sorted cells on Day 16. Increased proliferation was observed from the high secretors in both single-cell colonies and bulk-sorted populations. Significant difference claimed by one-way ANOVA with post-hoc Tukey Honestly Significant Difference (HSD) test (*$p<0.05$). D) EV production rate (EVs/cell) from single-cell colony and bulk-sorted populations quantified from conditioned media.

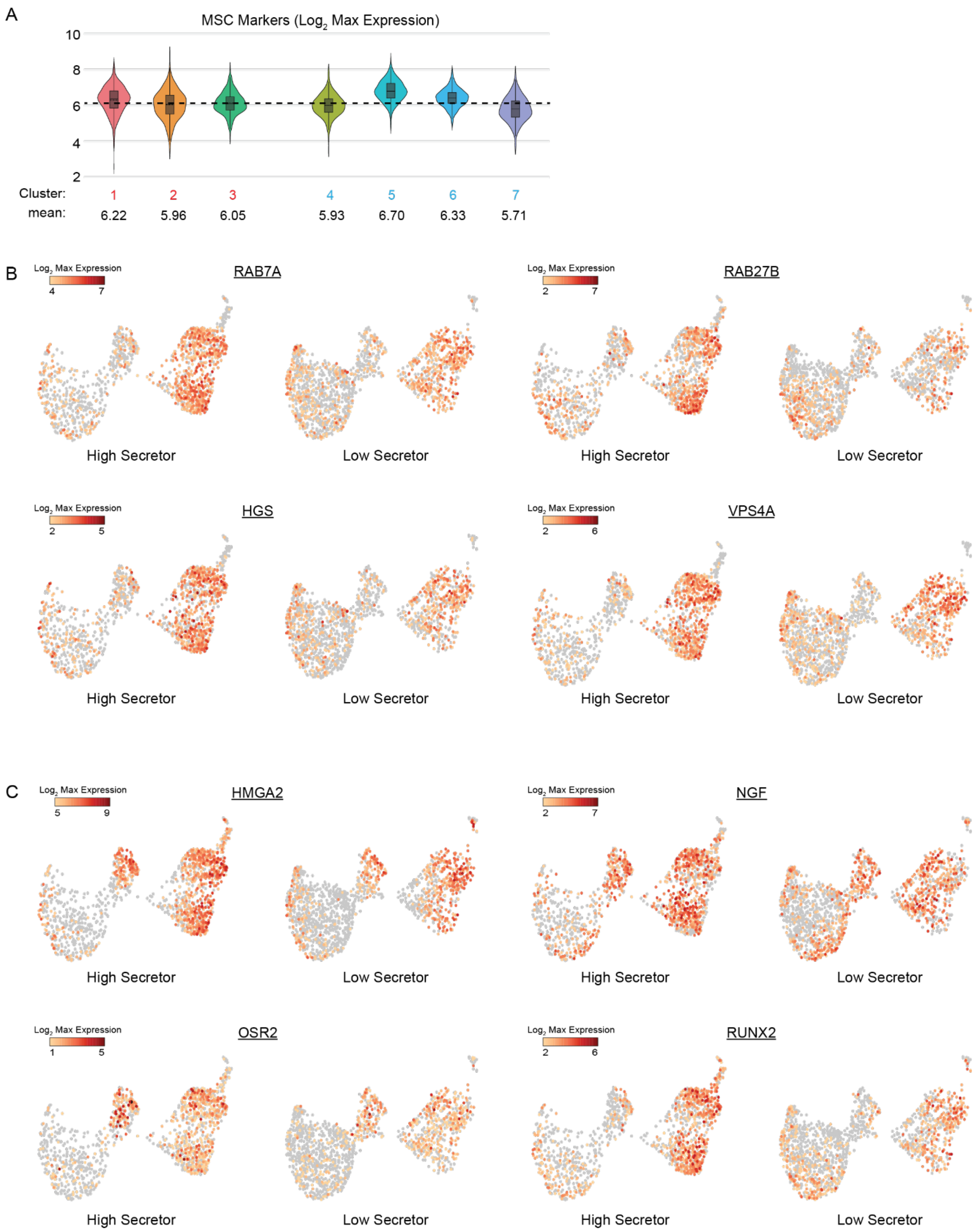


Supplementary Figure 4. Transcriptomic analysis of MSC marker and EV biogenesis expression. A) MSC marker expression was consistent within each cluster, indicating the nanovials do not affect intrinsic characteristic of stem cells. B) Genes associated with EV biogenesis (RAB7A, RAB27B, HGS, VPS4A) or C) with positive regulation of stem cell proliferation (HMGA2, NGF, OSR2, RUNX2) are specifically overexpressed among high secretors.

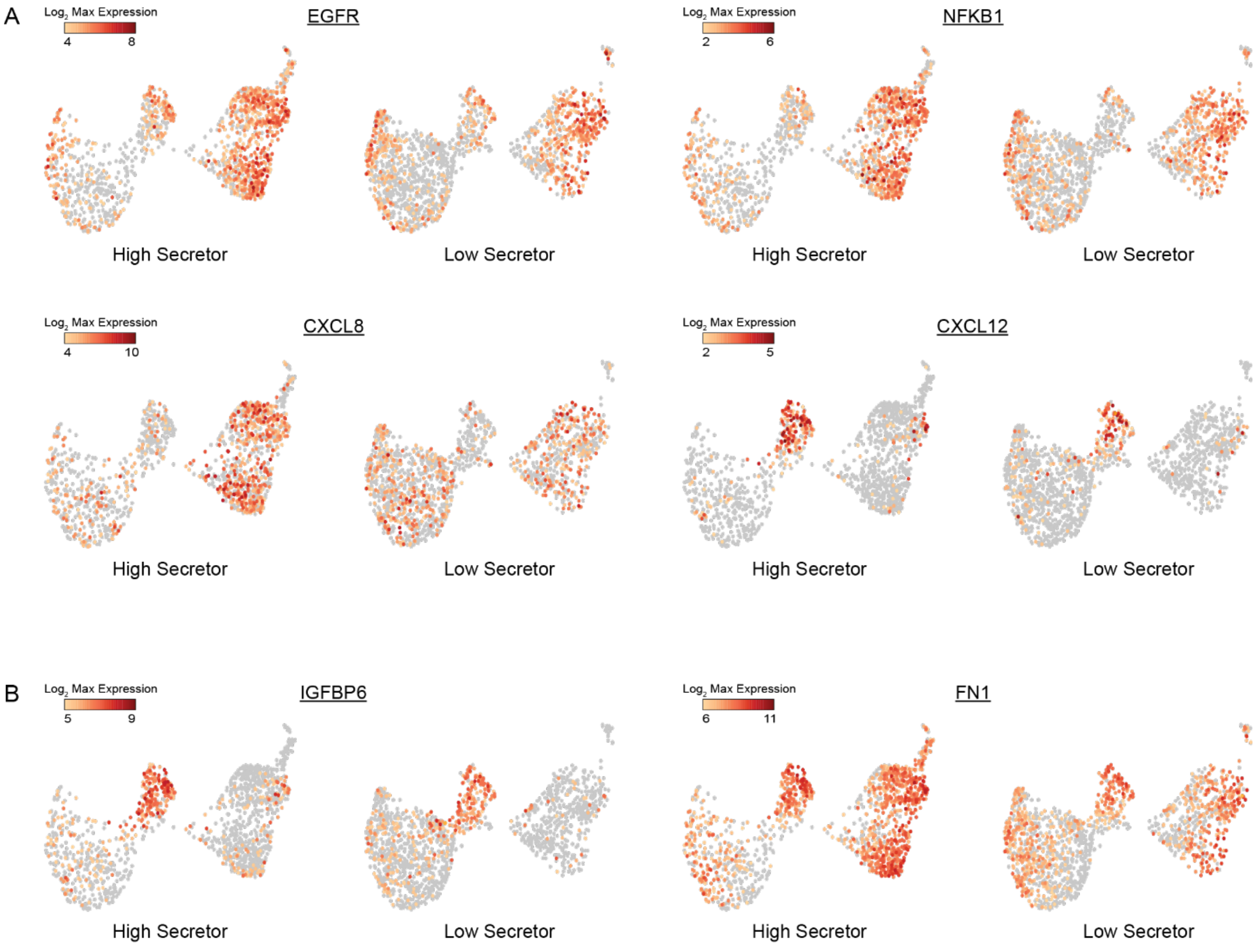


Supplementary Figure 5. Differences in expression level associated with tissue regeneration and vascular regenerative signal between high and low secretors. A) Pro-angiogenic factors such as EGFR, NFKB1, CXCL8, and CXCL12 are overexpressed among high secretors. B) Vascular regenerative signal associated genes such as IGFBP6 and FN1 are specifically expressed in cluster 3 and high secretors exhibited greater expression of these genes.

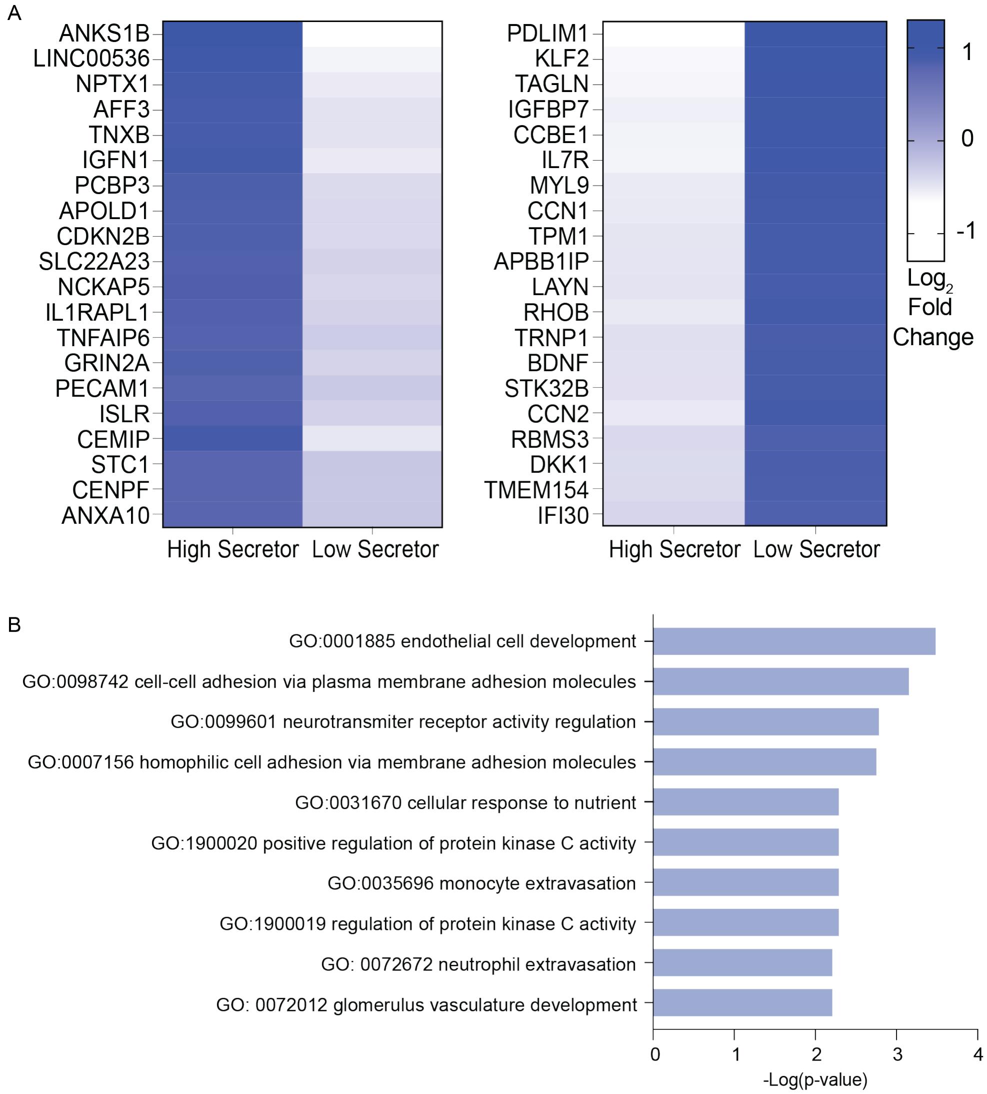


Supplementary Figure 6. Top 20 differentially expressed transcripts for high and low secretors and ontology terms associated with genes upregulated in high secretors. A) Distinct mRNA expression profiles in high vs. low EV secretors. The inclusion criteria of these transcripts was a 2-fold difference of log2 (fold-change) with a *p*-value <0.05. Blue signal, represents higher relative expression as compared to light blue signal. B) GO Biological Process annotations using the Enrichr toolkit for multiple testing with a *p*-value <0.01 (FDR<0.05) identified terms associated with the top 20 genes upregulated among high secretors.

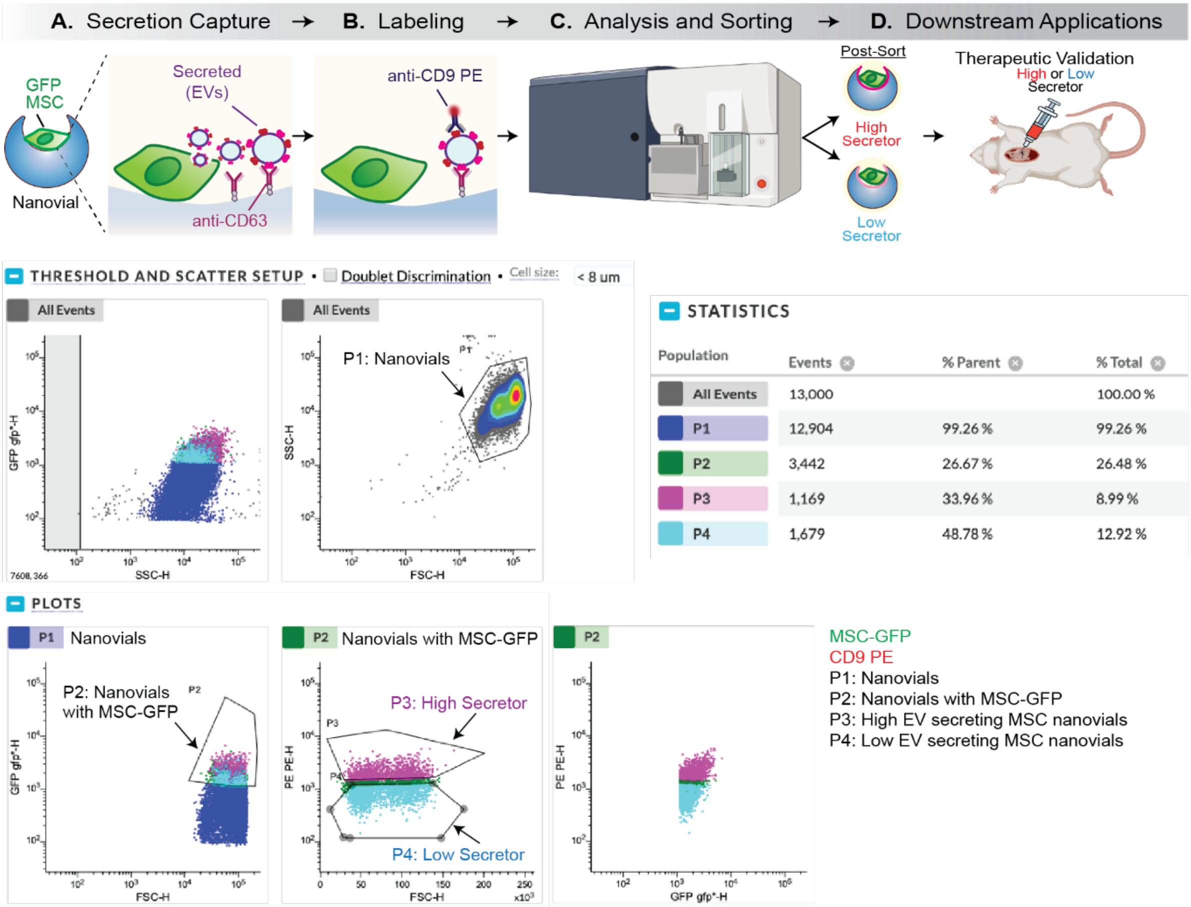


Supplementary Figure 7. Analysis and isolation of mouse MSCs based on EV secretion level. Schematic of the assay for mouse cells. Cells are loaded on anti-CD63 nanovials and sorted based on anti-CD9 PE signal along with cell marker (GFP).

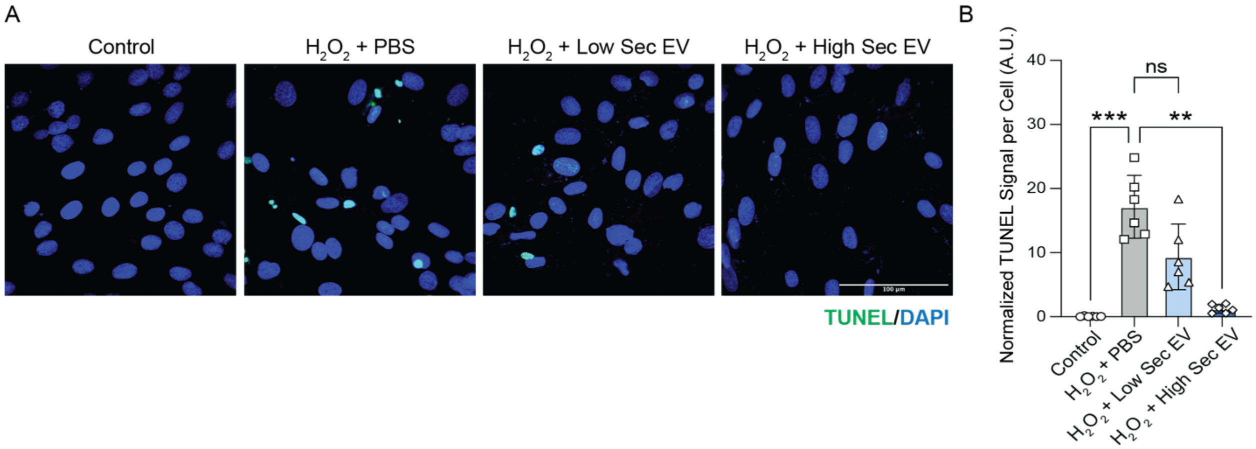


Supplementary Figure 8. EVs from high-sec MSCs exhibit a higher potential to reduce cell apoptosis following $H_2O_2$-induced rat cardiomyocyte (H9C2 cells) injury. A) TUNEL staining of cells. B) Quantification of apoptosis of cells (TUNEL+).

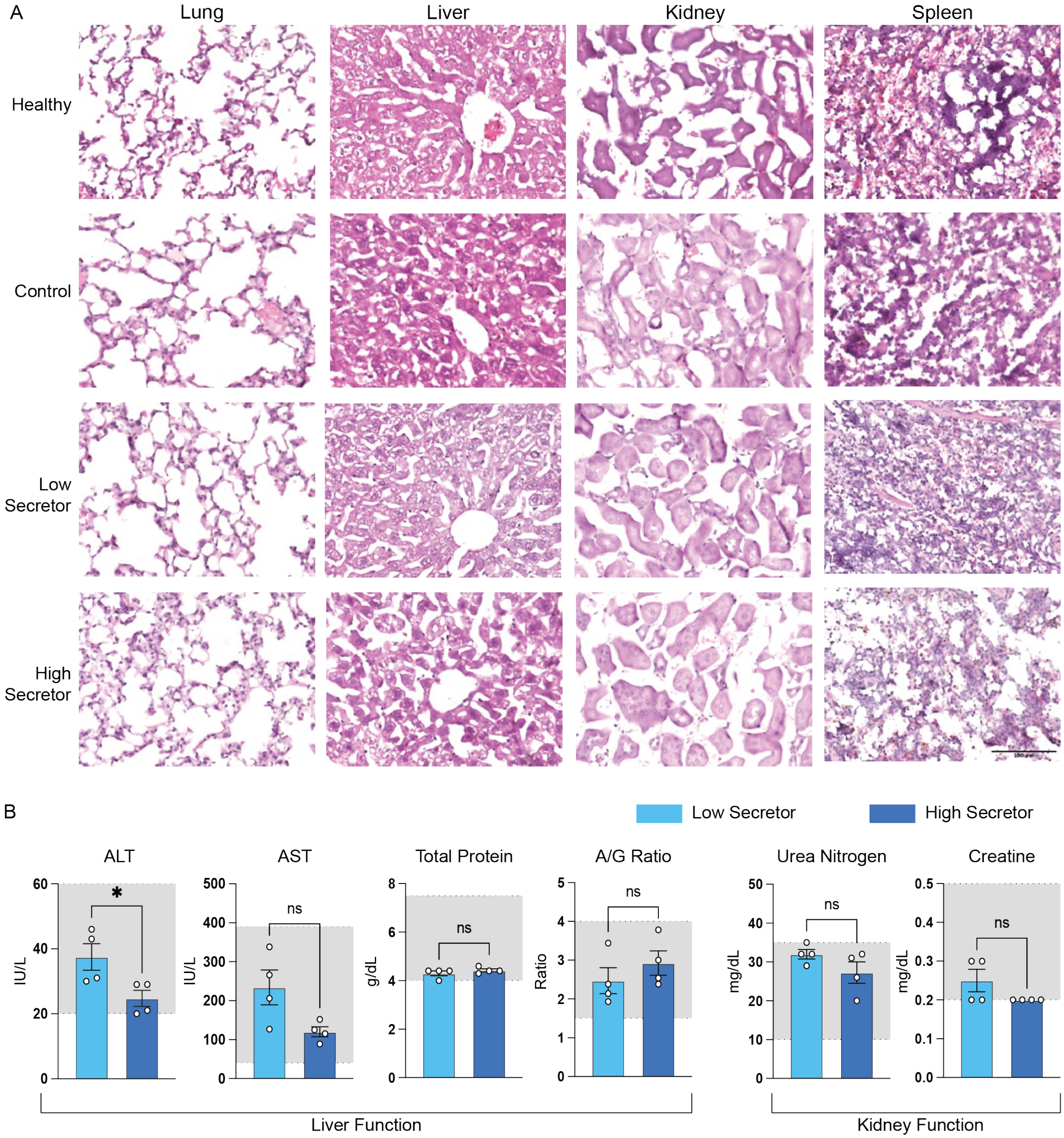


Supplementary Figure 9. Safety of high-sec and low-sec MSC treatment. (A) H&E staining of major organs (heart, lung, liver, spleen, kidney) from mice of different treatment groups. (B) Serum chemistry from mice of different treatment groups. (shaded region: range of normal values).
